# Renal outcomes of combination therapy with sodium-glucose cotransporter 2 inhibitors plus renin-angiotensin system blockers in patients with type 2 diabetes mellitus: A population-based cohort study

**DOI:** 10.1101/2024.08.28.610101

**Authors:** Ming-Hsien Tsai, Ming-chih Chen, Yen-Chun Huang, Wei-Shan Chang, Kai-Yuan Hsiao, Hung-Hsiang Liou, Yu-Wei Fang

## Abstract

**Background:** Sodium-glucose cotransporter 2 inhibitors (SGLT2i) can benefit patients with type 2 diabetes mellitus by reducing hazardous renal outcomes. This study aimed to evaluate whether the combination of sodium-glucose cotransporter 2 inhibitors (SGLT2i) and conventional renin-angiotensin system blockers (RASB) provides a synergistic effect on renal outcomes in patients with type 2 diabetes mellitus, compared to the combination of RASB and dipeptidyl peptidase 4 inhibitors (DPP4i).

**Methods:** This is a retrospective cohort study. The study utilized data from the Taiwan National Health Insurance Research Database (NHIRD), including patients with type 2 diabetes mellitus enrolled between January 1, 2016, and December 31, 2016. Participants were divided into two groups: the case group (n = 3,622) receiving RASB plus SGLT2i and the comparison group (n = 3,622) receiving RASB plus DPP4i. The groups were matched 1:1 based on gender, age, and Charlson comorbidity index. Additionally, TriNetX was used for external validation.

**Results:** Prior to matching, unadjusted hazard ratios (HRs) showed significant differences favoring the SGLT2i group for chronic kidney disease (CKD) (HR: 0.66; 95% CI, 0.58–0.74), advanced kidney failure (HR: 0.64; 95% CI, 0.44–0.93), and initiation of long-term dialysis (HR: 0.61; 95% CI, 0.38–0.97). These differences remained significant post-matching: CKD (HR: 0.74; 95% CI, 0.65–0.84), advanced kidney failure (HR: 0.62; 95% CI, 0.42–0.92), and commencement of long-term dialysis (HR: 0.53; 95% CI, 0.32–0.87). The renal benefits of the combination therapy were consistently observed in the TriNetX dataset.

**Limitations:** NHIRD lacks key clinical factors (e.g., physical features, lab data), potential baseline disparities due to retrospective design, and limited generalizability beyond Taiwanese patients, despite TriNexT validation.

**Conclusions:** In patients with type 2 diabetes mellitus, combination therapy with SGLT2i and RASB yielded better renal outcomes.

## Introduction

The prevalence of type 2 diabetes mellitus (T2DM) has increased over the past few decades in Taiwan [1]. Patients with T2DM carry a higher risk of cardiovascular disease (CVD), nephropathy, and stroke [2]. Furthermore, patients with T2DM without a history of CVD have an equal risk of cardiac events compared to those with a prior history of myocardial infarction[3]. This risk could be explained by the vascular damage from hyperglycemia, or one of several other existing comorbidities associated with increased risks, such as high blood pressure, dyslipidemia, and obesity. Diabetic kidney disease (DKD) is another complication of T2DM, seen in 25%–30% of patients with T2DM [4, 5] and accounting for one-third of end-stage renal disease cases requiring dialysis[6, 7].

Since the renin-angiotensin system (RAS) plays a major role in the renal pathophysiology of T2DM, its pharmacological inhibition by either angiotensin converting enzyme inhibitors (ACEIs) or angiotensin II receptor blockers (ARBs) can reduce cardiovascular (CV) mortality and all-cause mortality, as previously demonstrated [8–12]. As for DKD, using ACEIs and ARBs can have benefit in delaying the development and progression as revealed in several major randomized controlled trials[11, 13–15]. Therefore, the use of ACEIs and ARBs is considered the golden standard of management in high-risk of renal complications in T2DM.

Sodium-glucose cotransporter 2 inhibitors (SGLT2i) are new oral antidiabetic agents with a unique mechanism of action that functions independently of insulin secretion and action[16]. They lower plasma glucose by inhibiting glucose reabsorption in the renal proximal tube, producing glucosuria. This unique mechanism of action corrects several metabolic and hemodynamic abnormalities beyond glycemic control and reduces the incidence of nephropathy [17, 18].

Since both SGLT2i and RASB (i.e., ACEI or ARBs) demonstrate clinical benefit in T2DM, its combination use was inevitable, especially among patients at high risk. However, the real-world performance data supporting this practice is not sufficient [19–21]. This study aimed to analyze data from a nationwide Taiwanese database (real-world data) to shed light on the kidney safety and effectiveness of using SGLT2i together with RASB in treating patients with T2DM.

## Materials and Methods

### Study design and data sources

This research was a retrospective cohort study, drawing from a vast pool of information compiled within the National Health Insurance Research Database (NHIRD). This database is remarkably comprehensive, covering almost the entire population of Taiwan, a figure close to 99%, with records that have been meticulously collected since the year 1995. Prior to releasing the data analysis, Taiwan’s Ministry of Health, and Welfare (MOHW) adopted significant steps to ensure the privacy of the individuals whose data was included in our study. To this end, MOHW meticulously anonymized all the records of the beneficiaries making a claim, which effectively removed any personal identifiers from the data set.

Because of this careful anonymization, it was determined that there was no longer a necessity to obtain informed consent from the subjects of the study, since they could no longer be directly or indirectly identified. Additionally, we sought and subsequently received a formal exemption from the need for informed consent by the Institutional Review Board of the Shin-Kong Wu Ho-Su Memorial Hospital. This relief from the usual consent requirement came after careful consideration and was officially sanctioned, as evidenced by the IRB approval number provided, which is 20220712R for reference. The data for this study were accessed on Jaunary 9, 2023.

### Study population and exclusion criteria

This study analyzed a population of patients with T2DM, with an enrollment period from January 1, 2016, to December 31, 2016, based on the NHIRD (n = 1,937,938). (Fig 1) presents the flowchart outlining the process of patient selection. Patients in this study were required to have continuously used metformin and RASB medications for at least 3 months (n = 595,501). Under Taiwan’s national health insurance (NHI) rules, metformin is the main treatment for diabetes and recommended only for patients with a kidney function (eGFR) over 30 mL/min/1.73 m². This criterion enabled us to effectively exclude patients with severely compromised renal function (eGFR < 30 mL/min/1.73m²).

**Fig 1:**
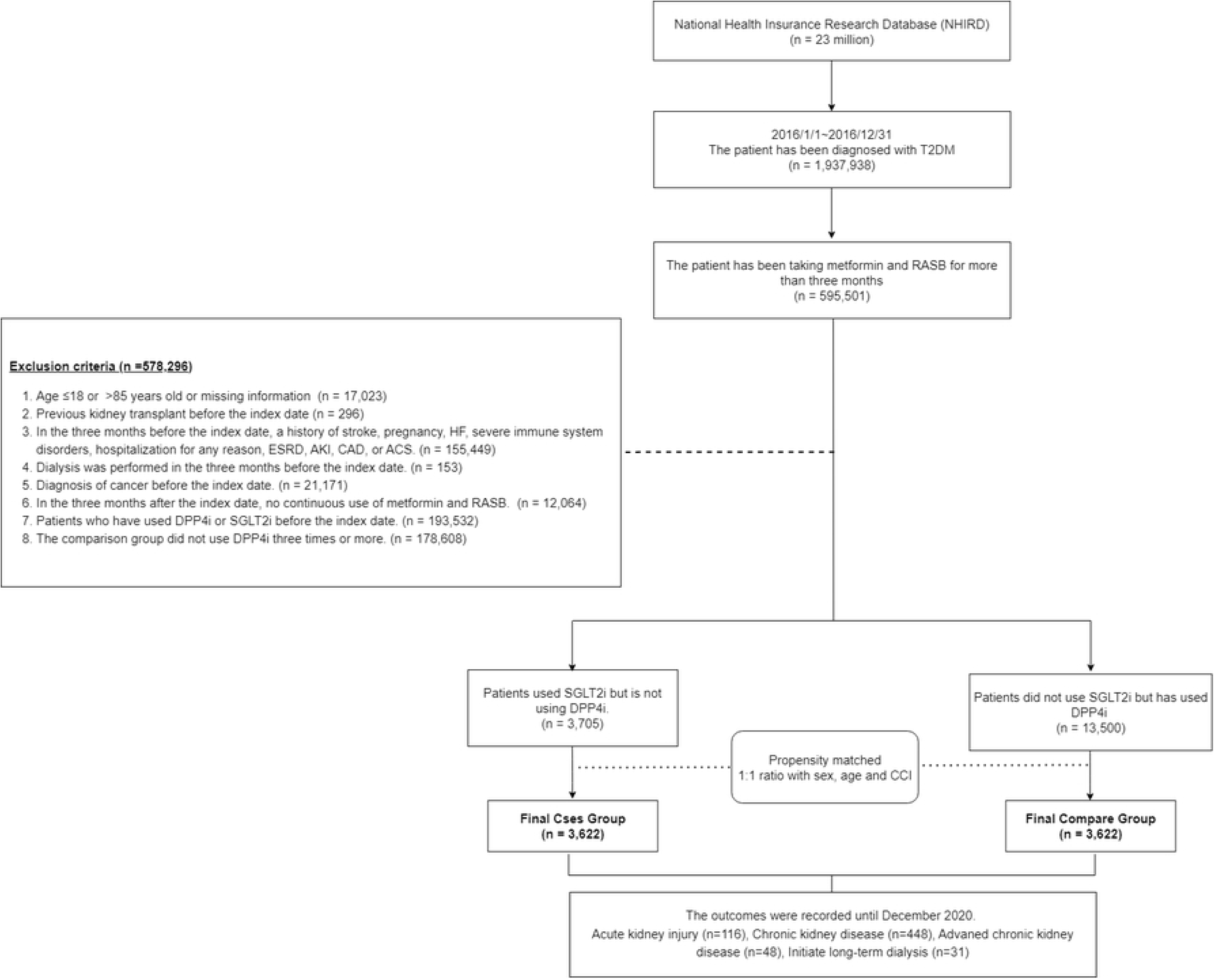
Schema of patient enrollment in the study Abbreviation: T2DM, type 2 diabetes mellitus; RASB, renin-angiotensin system blockers; HF, heart failure; ESRD, end-stage renal disease; AKI, acute kidney injury; CAD, coronary artery disease; ACS, acute coronary syndrome; DPP4i, dipeptidyl peptidase 4 inhibitors; SGLT2i, sodium–glucose cotransporter 2 inhibitors; CCI, Charlson comorbidity index.

The following were exclusion criteria for this study: (1) age less than 18 years or greater than 85 years or missing information (n = 17,023); (2) history of kidney transplant before the index date (n = 296); (3) history of stroke, pregnancy, heart failure (HF), immune system diseases, hospitalization for any reason, end-stage kidney disease (ESKD), acute kidney injury (AKI), coronary artery disease (CAD), or acute coronary syndrome (ACS) three months preceding the index date (n = 155,449); (4) history of dialysis three months prior to the index date (n = 153); (5) a diagnosis of cancer before the index date (n = 21,171); (6) discontinuation of metformin and RASB meds within three months from index date (n = 12,064); (7) use of either SGLT2i or DPP4i before the index date (n = 193,532); (8) patients in the comparison group who did not take DPP4i at least three times (n = 178,608).

The study population was divided into two groups. The case group included patients who used SGLT2i at least three times or more but had not used DPP4i (n = 3705). The comparison group consisted of patients who had not used SGLT2i but had used DPP4i three times or more (n = 13,500). The index date was defined as the day after three months of SGLT2i use, and the comparison group shared this index date. Patients were followed from the index date until the occurrence of an event, death, or the end of the database period (December 31, 2020). To reduce confounding variables, propensity score matching was done at a 1:1 rate, adjusting for gender, age, and Charlson Comorbidity Index (CCI). After matching, the case group comprised 3,622 individuals, mirroring the size of the comparison group.

### Covariates

The variables gathered included: gender, age, CCI score, and specific comorbidities (e.g., hypertension, ischemic heart disease, arrhythmia, atrial fibrillation, stroke, chronic obstructive pulmonary disease, asthma, peptic ulcer, dyslipidemia, gout, and liver cirrhosis). Past medication history was also reviewed, including beta-blockers, calcium channel blockers, alpha-blockers, anticoagulants, diuretics, antithrombotic, insulin, sulfonylureas, thiazolidinediones, acarbose, lipid-lowering agents, urate-lowering agents, non-steroidal anti-inflammatory drugs, and sedative-hypnotics. The disease and drug codes are shown in supplementary material (S1 Table).

### Study outcomes

This study commenced on the index date. Patients were followed until the occurrence of clinical events, including acute kidney injury (AKI), CKD, initiation of erythropoietin (ESA) therapy, and initiation of long-term dialysis. Under Taiwan’s national health insurance (NHI) rules, ESA payments are approved for CKD patients with an eGFR under 15 mL/min/1.73 m^2^ and a hemoglobin level below 9 g/dL. Thus, CKD patients getting ESA in the NHIRD are in stage 5 of CKD (eGFR <15 mL/min/1.73 m^2^). This stage 5 definition is widely used in research[22–24]. Other codes of outcome are shown in supplementary material (S1Table). If none of those clinical events occurred, this ongoing observation and data collection persisted until either the patient’s death or the end of the study period (December 31, 2020).

### External dataset validation

We tried to use a global dataset to validate our finding. TriNetX is the global federated health research network providing access to electronic medical records (diagnoses, procedures, medications, laboratory values, genomic information) across large healthcare organizations (HCOs). It allows users to search for patients meeting specific criteria in a de-identified database without the need for prior Institutional Review Board (IRB) approval [25]. This sensitive test was run on the set of HCOs grouped into a network called Global Collaborative Network, which this network included 119 HCOs.

The validation analysis was conducted on April 22, 2024. A population of individuals with diabetes, an eGFR of ≥ 30 mL/min/1.73 m², and the use of metformin and RASB was collected. From this group, two cohorts were established: one taking SGLT2 inhibitors (n = 41,411) and the other taking DPP4 inhibitors (n = 41,411), with each cohort matched on a propensity score. The analysis process is shown in the supplementary material (S2 Table).

### Statistical analysis

The demographic findings were shown as percentages or averages along with their standard deviations. We compared categories and continuous variables using the chi-square test and the t-test, respectively. To reduce differences between the case group and the comparison group in further analyses, we used a method called propensity score matching at a 1:1 ratio. Moreover, we used Cox proportional regression models to calculate hazard ratios (HRs) and 95% confidence intervals (CIs) to evaluate the risk of the outcomes. The proportional hazard assumption was assessed using Schoenfeld residuals.

The initial model was a crude analysis, whereas the subsequent model included further adjustments for all baseline confounding variables. Additionally, in the subgroup analysis, we included only the group with more than 500 patients. Two-tailed P-values less than 0.05 were considered statistically significant. All analytical procedures were performed using SAS version 9.4 (SAS Institute, Cary, NC, USA).

## Results

### Baseline characteristics

The average age of individuals using SGLT2i was 59.2 ± 12.1 years, and 56.7% were male. Among them, 91.8% had hypertension, 18.9% had a prior ischemic event, 9.9% had heart-related conditions, 19.4% had peptic ulcers, 17.1% had a history of stroke, and the average CCI score was 3.1 ± 2.0. Prior to matching, SGLT2i users had a higher prevalence of comorbidities and medication usage compared to DDP4i users. This difference between the groups persisted even after matching (Table 1).

**Table 1:**
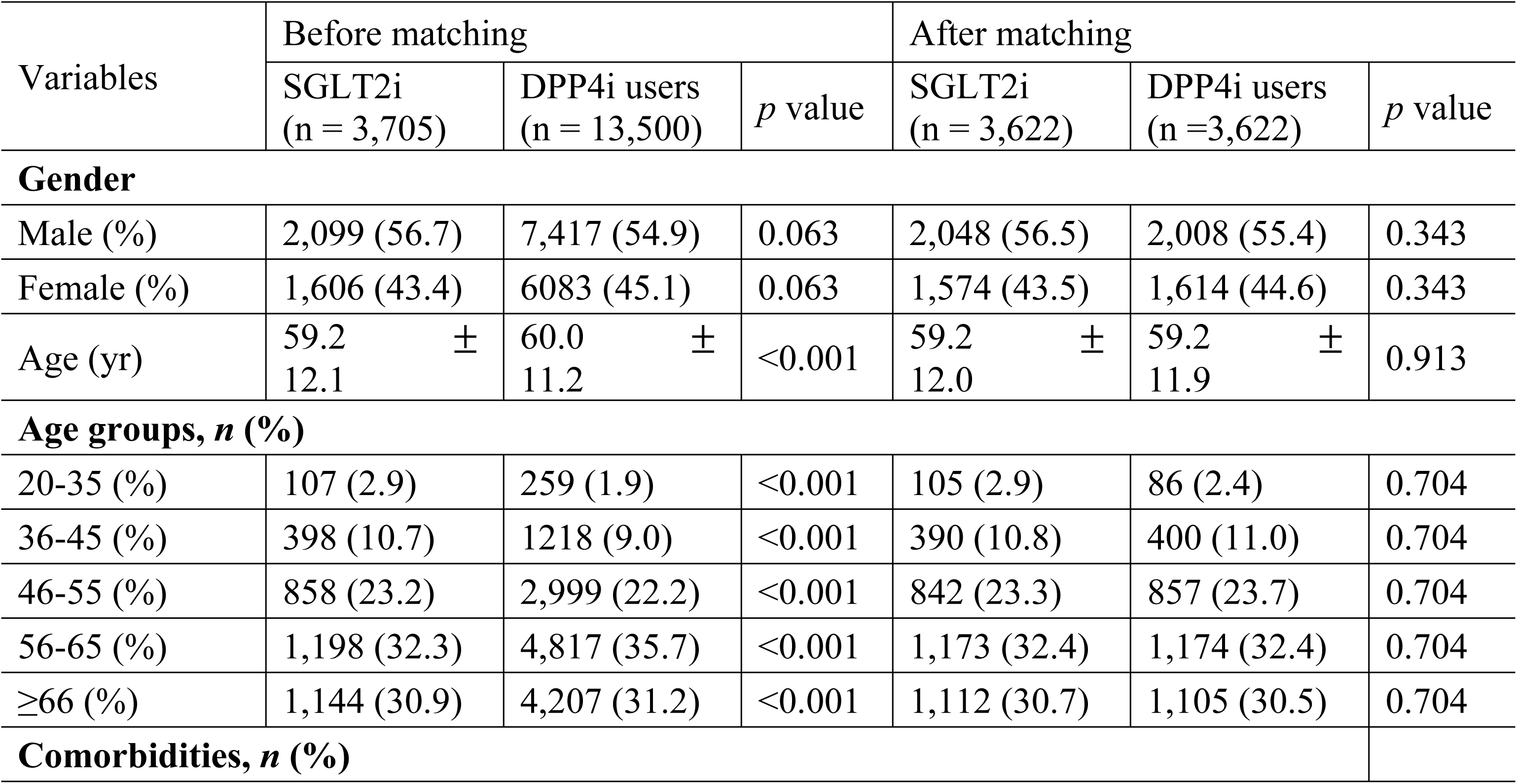
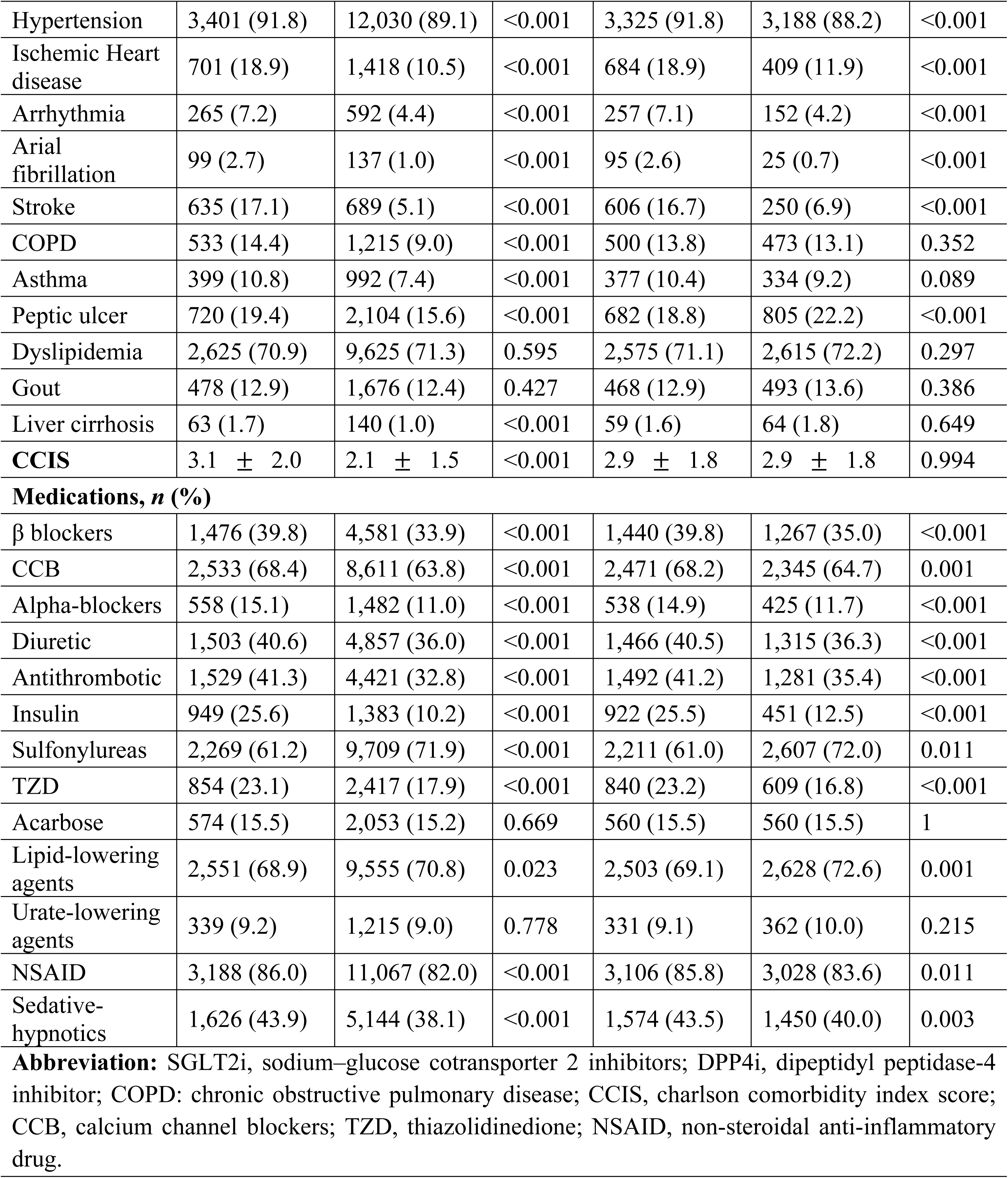
Demographic and clinical characteristics.

| Variables | Before matching |  |  | After matching |  |  |
| --- | --- | --- | --- | --- | --- | --- |
|  | SGLT2i<br>(n = 3,705) | DPP4i users<br>(n = 13,500) | *p* value | SGLT2i<br>(n = 3,622) | DPP4i users<br>(n = 3,622) | *p* value |
| **Gender** |  |  |  |  |  |  |
| Male (%) | 2,099 (56.7) | 7,417 (54.9) | 0.063 | 2,048 (56.5) | 2,008 (55.4) | 0.343 |
| Female (%) | 1,606 (43.4) | 6083 (45.1) | 0.063 | 1,574 (43.5) | 1,614 (44.6) | 0.343 |
| Age (yr) | 59.2 ± 12.1 | 60.0 ± 11.2 | <0.001 | 59.2 ± 12.0 | 59.2 ± 11.9 | 0.913 |
| **Age groups, *n* (%)** |  |  |  |  |  |  |
| 20-35 (%) | 107 (2.9) | 259 (1.9) | <0.001 | 105 (2.9) | 86 (2.4) | 0.704 |
| 36-45 (%) | 398 (10.7) | 1218 (9.0) | <0.001 | 390 (10.8) | 400 (11.0) | 0.704 |
| 46-55 (%) | 858 (23.2) | 2,999 (22.2) | <0.001 | 842 (23.3) | 857 (23.7) | 0.704 |
| 56-65 (%) | 1,198 (32.3) | 4,817 (35.7) | <0.001 | 1,173 (32.4) | 1,174 (32.4) | 0.704 |
| $\geq 66$ (%) | 1,144 (30.9) | 4,207 (31.2) | <0.001 | 1,112 (30.7) | 1,105 (30.5) | 0.704 |
| **Comorbidities, *n* (%)** |  |  |  |  |  |  |
| Hypertension | 3,401 (91.8) | 12,030 (89.1) | <0.001 | 3,325 (91.8) | 3,188 (88.2) | <0.001 |
| Ischemic Heart disease | 701 (18.9) | 1,418 (10.5) | <0.001 | 684 (18.9) | 409 (11.9) | <0.001 |
| Arrhythmia | 265 (7.2) | 592 (4.4) | <0.001 | 257 (7.1) | 152 (4.2) | <0.001 |
| Atrial fibrillation | 99 (2.7) | 137 (1.0) | <0.001 | 95 (2.6) | 25 (0.7) | <0.001 |
| Stroke | 635 (17.1) | 689 (5.1) | <0.001 | 606 (16.7) | 250 (6.9) | <0.001 |
| COPD | 533 (14.4) | 1,215 (9.0) | <0.001 | 500 (13.8) | 473 (13.1) | 0.352 |
| Asthma | 399 (10.8) | 992 (7.4) | <0.001 | 377 (10.4) | 334 (9.2) | 0.089 |
| Peptic ulcer | 720 (19.4) | 2,104 (15.6) | <0.001 | 682 (18.8) | 805 (22.2) | <0.001 |
| Dyslipidemia | 2,625 (70.9) | 9,625 (71.3) | 0.595 | 2,575 (71.1) | 2,615 (72.2) | 0.297 |
| Gout | 478 (12.9) | 1,676 (12.4) | 0.427 | 468 (12.9) | 493 (13.6) | 0.386 |
| Liver cirrhosis | 63 (1.7) | 140 (1.0) | <0.001 | 59 (1.6) | 64 (1.8) | 0.649 |
| **CCIS** | 3.1 ± 2.0 | 2.1 ± 1.5 | <0.001 | 2.9 ± 1.8 | 2.9 ± 1.8 | 0.994 |
| **Medications, *n* (%)** |  |  |  |  |  |  |
| β blockers | 1,476 (39.8) | 4,581 (33.9) | <0.001 | 1,440 (39.8) | 1,267 (35.0) | <0.001 |
| CCB | 2,533 (68.4) | 8,611 (63.8) | <0.001 | 2,471 (68.2) | 2,345 (64.7) | 0.001 |
| Alpha-blockers | 558 (15.1) | 1,482 (11.0) | <0.001 | 538 (14.9) | 425 (11.7) | <0.001 |
| Diuretic | 1,503 (40.6) | 4,857 (36.0) | <0.001 | 1,466 (40.5) | 1,315 (36.3) | <0.001 |
| Antithrombotic | 1,529 (41.3) | 4,421 (32.8) | <0.001 | 1,492 (41.2) | 1,281 (35.4) | <0.001 |
| Insulin | 949 (25.6) | 1,383 (10.2) | <0.001 | 922 (25.5) | 451 (12.5) | <0.001 |
| Sulfonylureas | 2,269 (61.2) | 9,709 (71.9) | <0.001 | 2,211 (61.0) | 2,607 (72.0) | 0.011 |
| TZD | 854 (23.1) | 2,417 (17.9) | <0.001 | 840 (23.2) | 609 (16.8) | <0.001 |
| Acarbose | 574 (15.5) | 2,053 (15.2) | 0.669 | 560 (15.5) | 560 (15.5) | 1 |
| Lipid-lowering agents | 2,551 (68.9) | 9,555 (70.8) | 0.023 | 2,503 (69.1) | 2,628 (72.6) | 0.001 |
| Urate-lowering agents | 339 (9.2) | 1,215 (9.0) | 0.778 | 331 (9.1) | 362 (10.0) | 0.215 |
| NSAID | 3,188 (86.0) | 11,067 (82.0) | <0.001 | 3,106 (85.8) | 3,028 (83.6) | 0.011 |
| Sedative-hypnotics | 1,626 (43.9) | 5,144 (38.1) | <0.001 | 1,574 (43.5) | 1,450 (40.0) | 0.003 |

**Abbreviation:** SGLT2i, sodium–glucose cotransporter 2 inhibitors; DPP4i, dipeptidyl peptidase-4 inhibitor; COPD: chronic obstructive pulmonary disease; CCIS, charlson comorbidity index score; CCB, calcium channel blockers; TZD, thiazolidinedione; NSAID, non-steroidal anti-inflammatory drug.

### The hazardous kidney outcome in the combination of SGLT2i with RASB

Fig 2A illustrates the Kaplan-Meier survival curves for AKI among groups using SGLT2i and DPP4i. The survival difference between these two groups was not statistically significant for AKI, as determined by a log-rank test with a p-value of 0.125. Table 2 shows the rates of AKI in patients using SGLT2i and DPP4i after they have been matched, with figures of 0.8 and 0.6 per 100 person-years, respectively. The likelihood of experiencing acute renal failure as a side effect was comparable between the SGLT2igroup and the DPP4i group. The risk difference was statistically insignificant in both the preliminary analysis (HR, 1.24; 95% CI, 0.94–1.63; p=0.126) and the multivariable analysis that adjusted for other factors (HR, 1.16; 95% CI, 0.87–1.55; p=0.304).

**Fig 2:**
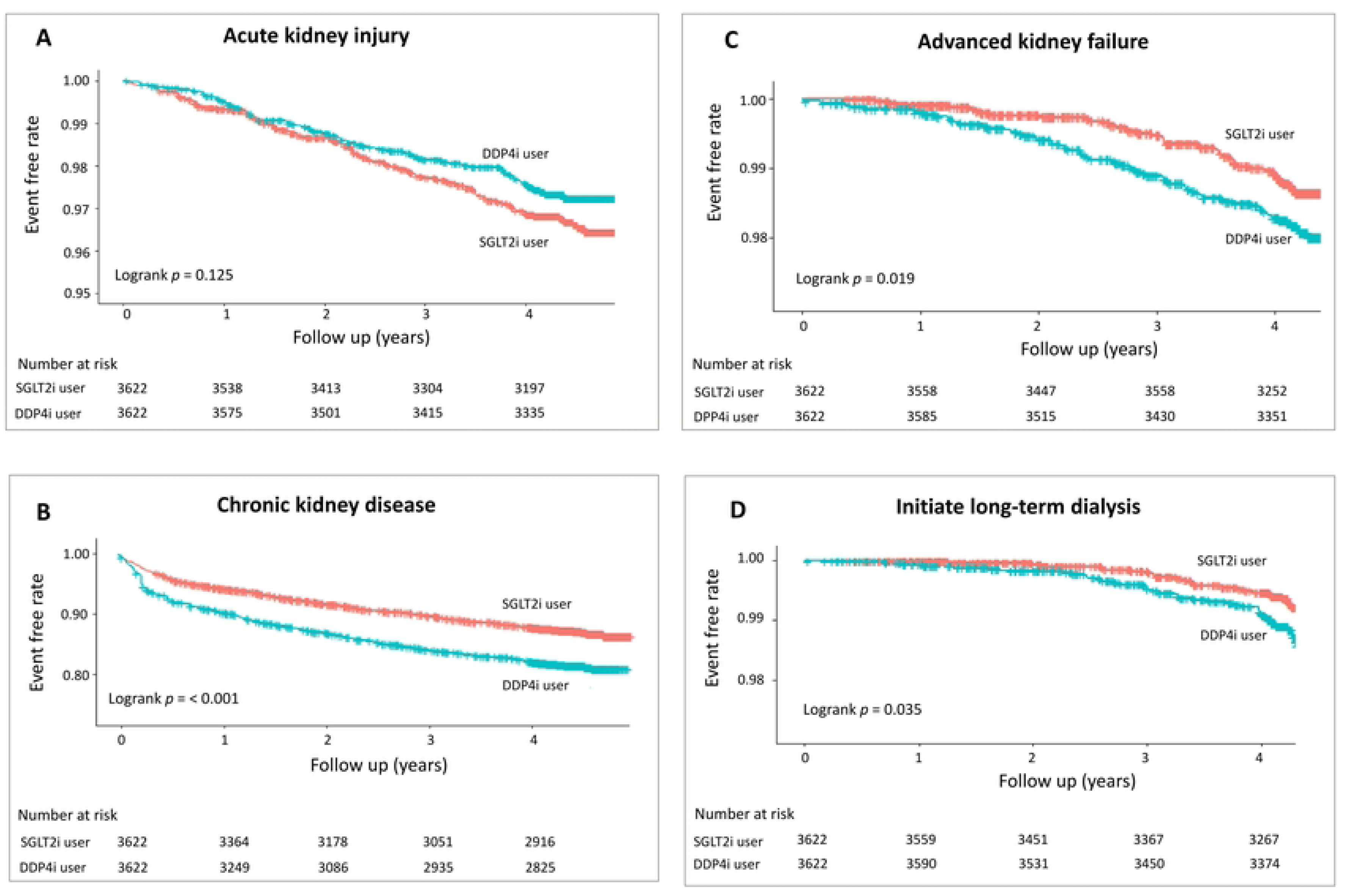
Kaplan–Meier cumulative event-free plots of (A) acute kidney injury, (B) chronic kidney disease, (C) advanced kidney failure, and (D) initialization of long-term dialysis. Abbreviation: SGLT2i, sodium–glucose cotransporter 2 inhibitors; DDP4i, dipeptidyl peptidase-4 inhibitor.

**Table 2.**
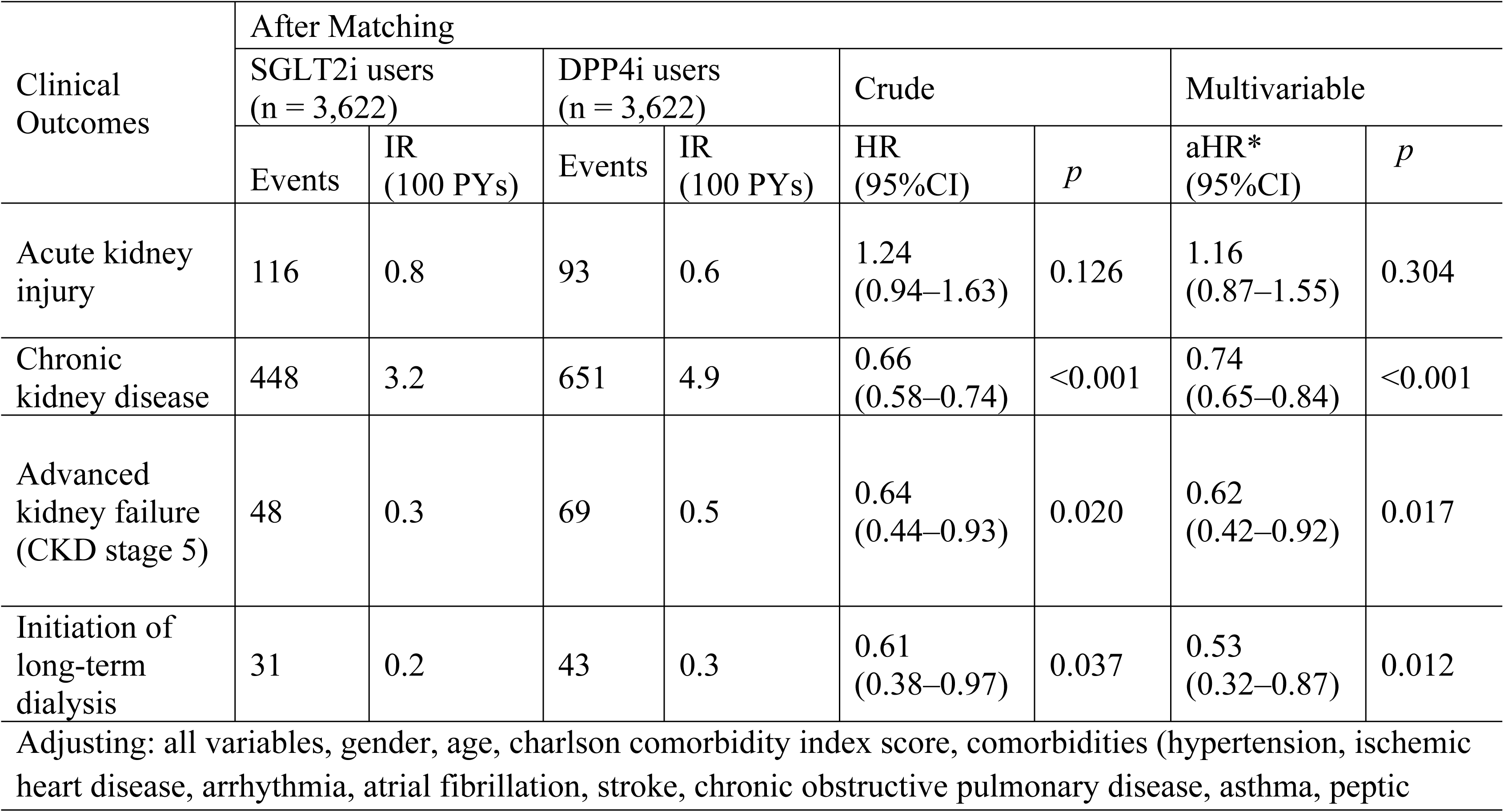
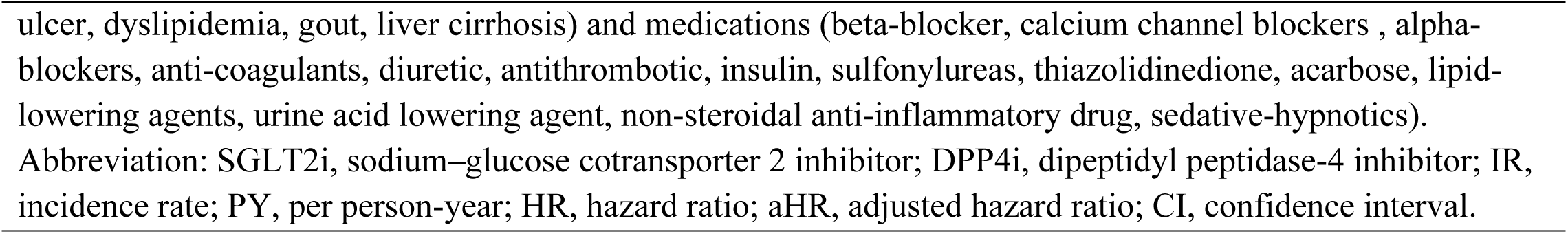
Risk of clinical outcomes in diabetic patients with RASB comparing SGLT2i users vs. rs. DDP4i users.

### The benefits of combining SGLT2i and RASB on Kidney Health

Table 2 presents the incidence rates of several clinical events among SGLT2i users and DDP4i users in patients after matching, including CKD (3.2 vs. 4.9 per 100 person-year), advanced kidney failure (0.3 vs. 0.5 per 100 person-year), and initiation of long-term dialysis (0.2 vs. 0.3 per 100person-year). The differences of Kaplan-Meier survival curves between groups taking SGLT2i and DPP4i were statistically significant for CKD (p <0.001), advanced kidney failure (p = 0.019) and initiation of long-term dialysis (p = 0.035) (Fig 2B–D).

Moreover, the unadjusted HR was significant for CKD (0.66; 95% CI, 0.58–0.74, p < 0.001), advanced kidney failure (0.64; 95% CI, 0.44–0.93, p < 0.020), and initiation of long-term dialysis (0.61; 95% CI, 0.38–0.97, p < 0.037). After matching, these three kidney outcomes still showed significant differences: the HR for CKD was 0.74 (95% CI, 0.65–0.84, p < 0.001), for advanced kidney failure was 0.62 (95% CI, 0.42–0.92, p < 0.017), and for beginning long-term dialysis was 0.53 (95% CI, 0.32–0.87, p < 0.012) (Table 2).

### Subgroup analysis

A series of stratified analyses were conducted to evaluate the reliability of our findings in (Fig 3). The reduced HRs of CKD development among diabetic patients with RABS use in favor of SGLT2i use were consistent across almost all patient subgroups, except those with peptic ulcer and those without lipid-lowering agents (Fig 3A). In advanced kidney failure, only half of the subgroups experienced benefits from using a combination of SGLT2i and RASB. Nevertheless, the data suggests a favorable trend adopting this combination therapy (Fig 3B).

**Fig 3:**
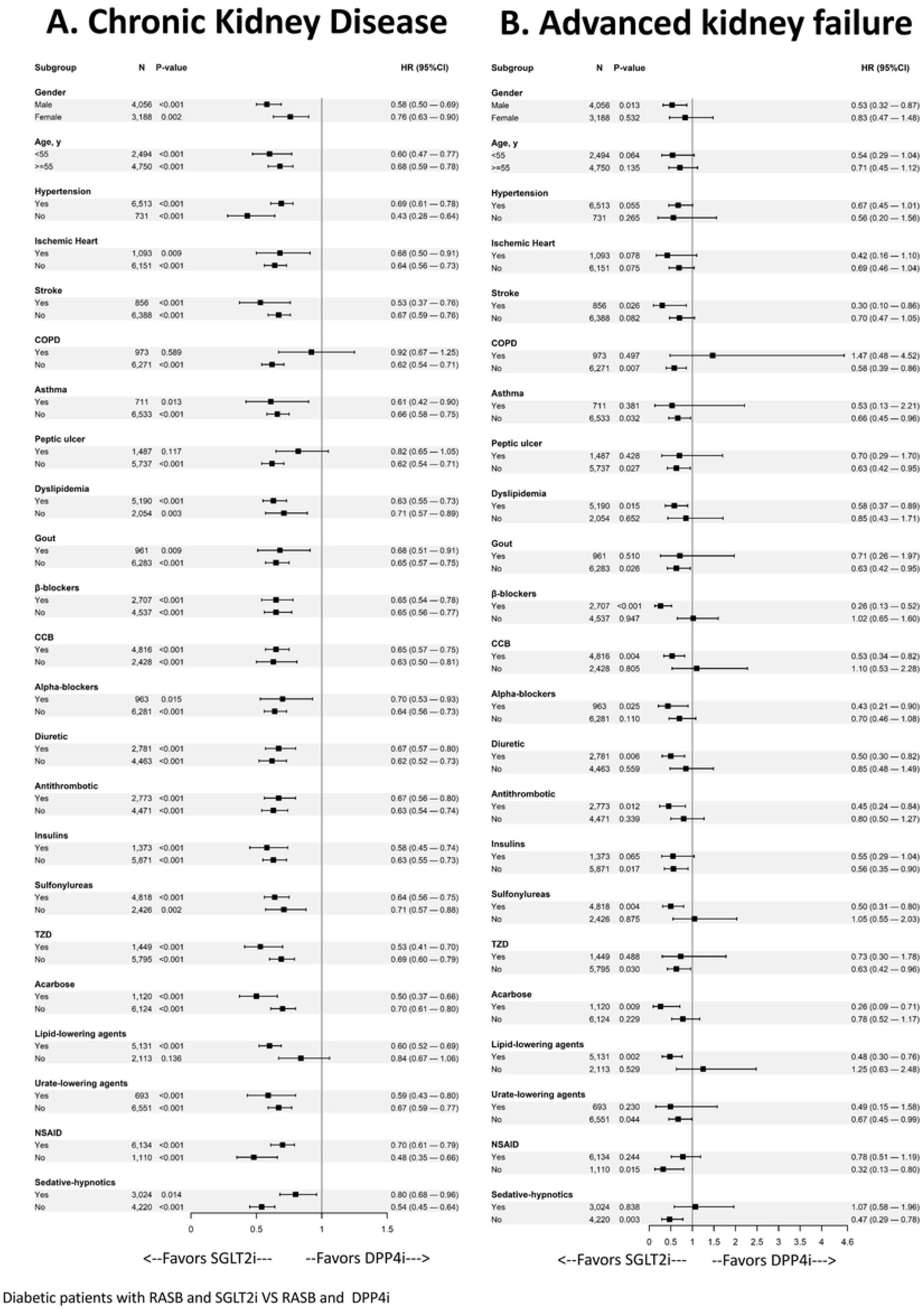
Subgroup analysis of the effect of combining SGLT2i and RASB on the occurrence of chronic kidney disease and advanced kidney failure Abbreviation: CCB, calcium channel blockers; RASB, renin-angiotensin system blockers; SGLT2i, sodium–glucose cotransporter 2 inhibitors; DDP4i, dipeptidyl peptidase-4 inhibitor.

### External data validation

The table 3 shows that RASB and SGLT2i users had significantly lower rates of acute kidney injury (15.2% vs. 22.5%), advanced kidney failure (0.8% vs. 2.0%), and end stage kidney disease (4.1% vs. 3.3%) compared to RASB and DPP4i users, with HRs ranging from 0.63 to 0.91 (p <0.001 for all outcomes). Such findings were compatible with our finding, mentioned above.

**Table 3.** External validation of the risk of kidney outcomes in diabetic patients using renin-angiotensin system blockers with TriNetX dataset.

| Clinical Outcomes | After Matching |  |  |  |  |  |
| --- | --- | --- | --- | --- | --- | --- |
|  | SGLT2i users<br>(n = 41,411) |  | DPP4i users<br>(n = 41,411) |  | SGLT2i users vs. DPP4i<br>users |  |
|  | Events (n) | (%) | Events(n) | (%) | HR (95%CI) | *p* |
| Acute kidney injury | 6,281 | 15.2 | 9,314 | 22.5 | 0.91 (0.88–0.93) | <0.001 |
| Advanced kidney failure | 332 | 0.8 | 810 | 2.0 | 0.63 (0.55–0.72) | <0.001 |
| End stage of kidney disease | 1,701 | 4.1 | 2,778 | 6.7 | 0.79 (0.74–0.84) | <0.001 |

Propensity score matching by age, gender, comorbidities (hypertension, ischemic heart disease, Cerebrovascular diseases, atrial fibrillation, asthma, chronic obstructive pulmonary disease, dyslipidemia, gout, liver cirrhosis, peptic ulcer) and medications (beta-blocker, calcium channel blockers, alpha-blockers, diuretic, insulin, lipid-lowering agents, urine acid lowering, and pioglitazone).
Abbreviation: SGLT2i, sodium–glucose cotransporter 2 inhibitor; DPP4i, dipeptidyl peptidase-4 inhibitor; HR, hazard ratio; CI, confidence interval.

## Discussion

This population-based cohort study demonstrated that combination use of SGLT2i plus RASB has clinical benefits in participants with early T2DM who used metformin. This combination demonstrated improvement in renal outcomes, including the prevention of CKD development, delaying the progression to severe kidney failure (identified as CKD stage 5), and averting end-stage kidney disease. The study also found no increased risk of AKI with this combination treatment compared to the use of DPP4i and RASB. Additionally, subgroup analyses indicate a positive trend in favor of adopting combination therapy. Thus, the evidence suggests that the combination of SGLT2i and RASB may be more effective in promoting kidney health in T2DM patients.

SGLT2i have emerged as a promising drug class in clinical practice. Traditionally, SGLT2i are reserved for adjunctive therapy in T2DM if adequate control cannot be achieved by metformin. However, SGLT2i can also be used as monotherapy in cases wherein metformin is contraindicated[26]. Beyond their role in glucose control, SGLT2i have unique mechanisms that can benefit individuals with CVD, such as reduced mortality, lower risk of hospitalizations due to heart failure, enhanced blood pressure regulation, and improved overall cardiovascular health[17, 27–29]. In a meta-analysis of several clinical randomized controlled trials (EMPAREG OUTCOME, CANVAS, and DECLARE-TIMI 58) by Zelniker et al., T2DM patients with previous atherosclerotic CVD can benefit from SGLT2i, with improved cardiovascular outcomes such as new-onset or recurrent heart failure and hospitalization[30].

Renoprotective effects have also been reported for SGLT2i, with reduced composite renal outcomes, such as serum creatinine doubling, macroalbuminuria, and end-stage kidney disease in previous large-scale studies (EMPA-REG, DECLARE, and CANVAS) [18, 29, 31]. In a meta-analysis of those trials, SGLT2i reduced the incidence of worsening renal function, end-stage renal disease or renal death by 45% (HR 0.55; 95% CI 0.48–0.64) [30]. Other studies investigated the renoprotective effects of CKD, including CREDENCE for DKD and DAPA-CKD for patients with or without T2DM[32, 33]. Both of those trials found that SGLT2i can reduce proteinuria and delay CKD progression into end-stage kidney disease in diabetic and non-diabetic patients alike.

The use of ACEIs or ARBs is considered the gold standard for slowing down DKD development and progression, as backed by positive results in previous randomized controlled trials (RCTs) [11, 13–15]. Angiotensin II (Ang II), as a key mediator in the pathogenesis of DKD, can constrict the efferent arterioles to increase intraglomerular pressure, resulting in glomerular damage and proteinuria [34]. Ang II also participates in renal inflammation and fibrosis through NADPH oxidase to induce excess oxidative stress [34]. Therefore, RAAS blockade can vasodilate the efferent glomerular arterioles and lower intraglomerular pressure [11].

SGLT2i exerts its glucose-lowering effects by inhibiting sodium and glucose reabsorption in the kidney. Along with glycosuria, increasing sodium delivery to the macula densa induced tubuloglomerular feedback, thus promoting afferent vasoconstriction and efferent vasodilation as demonstrated in a micropuncture study [35, 36]. However, there is still controversy regarding the synergistic effects of combination therapy with RAAS blockade plus SGLT2i, since intraglomerular pressure dipping may contribute to further eGFR reduction and increase risk of AKI. In our cohort, combination therapy reduced the risk of AKI as well as the rate of advanced kidney failure and ESRD compared to controls.

Since SGLT2i has a unique role in treating patients with T2DM, the efficacy of its combination with ACEI/ARBs has been a crucial issue. Lytvyn Y et al. conducted a prospective study to evaluate the effects of combination therapy in type 1 DM patients with renal hyperfiltration [21]. Adding SGLT2i to RASB resulted in an initial drop in GFR as expected, but there was a benefit of reduced oxidative stress and improved blood pressure as well. Recent systematic reviews and meta-analyses have also confirmed that combination therapy can significantly reduce albuminuria, but with no difference in eGFR compared to standard care[19, 20]. Regarding outcomes, Zhao LM et. al. demonstrated that combination therapy had greater efficacy in lowering MACE, cardiovascular death or HHF, and composite kidney outcomes compared to ACEI/ARB monotherapy among T2DM patients[37]. These results were consistent with our real-world database analysis.

In our cohort, combination therapy reduced the incidence of CKD in most subgroups. Among patients with advanced kidney failure, those with female sex, underlying gout, asthma, and peptic ulcer disease seemed to have less benefit from combination therapy. Patients with history of NSAID use also did not benefit from combination therapy, likely due to its nephrotoxicity. NSAIDs can induce AKI by interfering with the production of prostaglandins, which are key vasodilators within kidneys. Decreased prostaglandin levels could lead to reduced renal perfusion, leading to ischemia especially under the setting of RAAS inhibition [38].

Our study had several strengths. First, a nationwide database was used, ensuring broad relevance for our findings. This finding was also validated by global dataset (TriNetX) Second, we specifically selected T2DM individuals using metformin, ensuring similar CKD stages in both case and comparison groups, (eGFR > 30 mL/min/1.73m^2^). However, this study also has some potential limitations. First, key confounding factors which could be clinically relevant, such as physical features, lifestyle, and laboratory data were not included in the NHIRD, and thus we were unable to consider those in our analysis. Second, since this was a retrospective cohort analysis and not an RCT, there could be baseline disparities between groups, potentially introducing bias due to confounding factors. To minimize potential bias, however, propensity score matching and multivariable adjustments in the regression model were performed. Third, since the study focused solely on Taiwanese patients, caution should be taken in applying these findings to broader global populations. However, the global dataset (TriNexT) validated our finding.

## Conclusion

Individuals with type 2 diabetes could see better kidney health outcomes by combining SGLT2i with RASB in real-world settings. This combination therapy is recommended for patients with diabetes and CKD to preserve kidney function.

## Author Contributions

**Conceptualization:** Hung-Hsiang Liou, Ming-Hsien Tsai, Yu-Wei Fang

**Formal analysis:** Ming-Chih Chen, Yen-Chun Huang, Wei-Shan Chang, Kai-Yuan Hsiao

**Supervision:** Ming-Chih Chen, Yu-Wei Fang

**Validation:** Ming-Hsien Tsai, Ming-Chih Chen

**Writing – original draft:** Ming-Hsien Tsai, Yu-Wei Fang

**Writing – review & editing:** all authors.

## Acknowledgements

We are grateful to the Health and Welfare Data Science Center (HWDC) and TriNetX for providing the data used in this study.

## Data availability statement

The dataset used in this study is held by the Taiwan Ministry of Health and Welfare (MOHW). Owing to the General Data Protection Regulation, the dataset is not available on request from the corresponding author. Any researcher interested in accessing this dataset can apply for access. Please visit the website of the National Health Informatics Project of the MOHW (https://dep.mohw.gov.tw/DOS/mp-113.html). TriNetX is a network connecting multiple research centers that offers instant access to anonymized data from the electronic health records of participating health-care organizations. Researchers can access this database at https://live.trinetx.com

## Supporting information

**S1 Table.** Code for comorbidities & drug. (PDF)

**S2 Table.** The cohort’s creation and outcome analysis in TriNetX (PDF)

